# Harnessing Tumor-Specific Transcript Diversity Uncovers a Shared Neoantigen Reservoir for Pancreatic Ductal Adenocarcinoma

**DOI:** 10.64898/2026.02.10.705024

**Authors:** Jingjing Zhao, Qiaojuan Li, Peng Lin, Yu Yang, Hongwu Yu, Yifan Wen, Wenqian Yu, Huiyi He, Sichen Tao, Feifei Zhang, Yan Li, Zhixiang Hu, Jing Xie, Zhen Chen, Shenglin Huang

**Author notes:** These authors contributed equally to this work (Jingjing Zhao, Qiaojuan Li, Peng Lin). **Corresponding author**: Shenglin Huang, Zhen Chen, and Jing Xie.

## Abstract

Pancreatic ductal adenocarcinoma (PDAC) is refractory to immunotherapy due to its immunologically cold microenvironment and the scarcity of mutation-derived neoantigens. Here, we introduce NeoAPP, a computational tool designed to systematically decode neoantigens arising from tumor-specific transcripts (TSTs) generated by transcriptional dysregulation. Multi-cohort transcriptomic profiling of 413 PDAC samples using NeoAPP reveal a median of 351 neoantigens per sample derived from 56 neoantigen-encoding TSTs (neoTSTs), surpassing mutation-derived counterparts in both abundance and patient coverage. Mechanistic analyses show that non-canonical splicing junction and transposable element activation drive neoantigen generation, while FOXA2-regulated promoter usage constitutes a potential major source of neoTSTs. Tumor-derived neoTSTs are also detected in extracellular vesicles and cancer-associated fibroblasts, implicating stromal crosstalk in immune modulation. Vaccination with neoTSTs induce CD8+ T cell responses in HLA-A*02:01/A*11:01 transgenic mice and suppressed tumor growth in syngeneic PDAC models. Collectively, this work establishes TST-derived neoantigens as a dominant and therapeutically actionable antigen reservoir in PDAC, advancing a transcriptome-guided framework for neoantigen discovery with potential to overcome immune resistance in low-mutation cancers.

**Abstract Figure:** 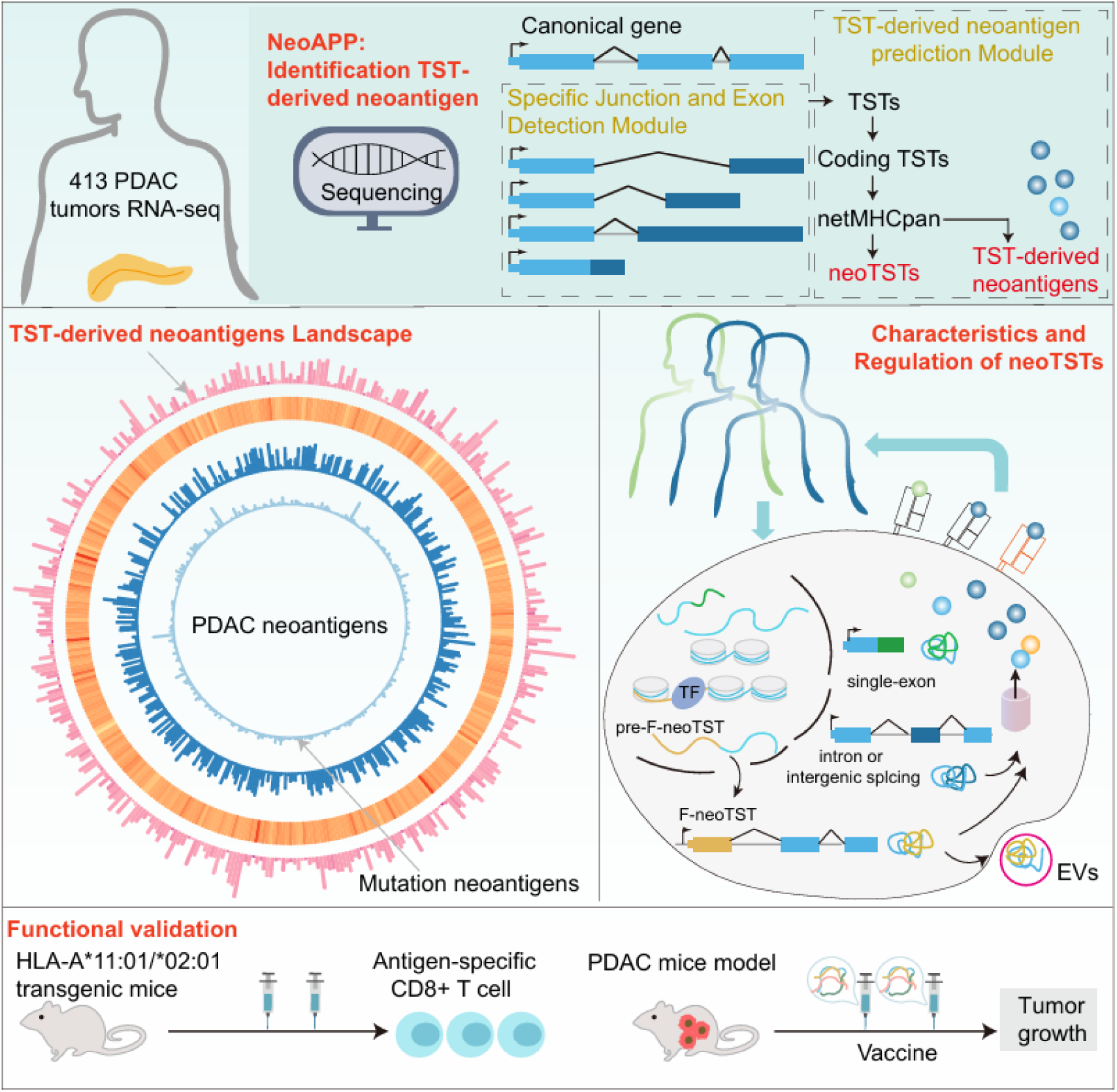

## 1 Introduction

Pancreatic ductal adenocarcinoma (PDAC) remains among the most lethal solid tumors, accounting for over 5% of global cancer mortality while sustaining a 5-year survival rate of 12%. While first-line regimens such as FOLFIRINOX demonstrate limited clinical benefit in advanced PDAC^[1, 2]^, these marginal gains underscore the imperative for paradigm-shifting therapeutic approaches.

The renaissance of neoantigen-targeted immunotherapy has revolutionized cancer treatment, demonstrating unprecedented clinical responses across multiple tumor types. Personalized mutation-derived neoantigen vaccines have shown durable clinical benefit in melanoma and NSCLC through robust T cell activation^[3, 4]^, while analogous strategies are transforming therapeutic paradigms for renal cell carcinoma (RCC) and hepatocellular carcinoma (HCC)^[5–7]^. Recent breakthrough work revealed that combinatorial adjuvant therapy with individualized mRNA neoantigen vaccines and checkpoint inhibition in resected PDAC patients induced de novo high-magnitude neoantigen-specific T cell responses^[8, 9]^, effectively preventing recurrence in this traditionally immunologically “cold” malignancy. Despite these advances, only ∼50% of PDAC patients exhibit vaccine responsiveness^[8]^, a limitation attributed to the malignancy’s low tumor mutation burden (TMB) compounded by its intrinsic genomic stability. Over 98% of PDAC cases exhibit microsatellite stability (MSS)^[10, 11]^, a hallmark of immunologically “cold” tumors with sparse immunogenic mutations. Systematic analyses reveal that merely 1.6% of somatic mutations yield immunogenic neoantigens, 99% of which are patient-specific and non-shared between individuals^[12,13]^. These dual constraints, low TMB-driven antigen paucity and the predominance of private neoantigens, create an insurmountable bottleneck in neoantigen discovery pipelines for PDAC, necessitating alternative strategies to expand the targetable immunopeptidome.

Transcriptional splicing aberrations represent an underutilized reservoir of tumor-specific antigens (TSAs) with unique advantages in stability and predictability. For instance, Kwok et al. demonstrated that GNAS neojunctions serve as HLA-restricted immunogens in glioma^[14]^. Several studies also have characterized splicing-derived epitopes originating from transposable elements (TEs) and noncoding regions, revealing non-canonical ORFs that expand the immunogenic landscape across malignancies^[15–18]^. While multiple computational approaches exist for identifying splicing junctions, our ASJA algorithm^[19]^ demonstrates superior accuracy in detecting both annotated and novel splicing events^[20]^. Leveraging this platform, we successfully identified oncogenic tumor-specific transcripts (TSTs)^[21]^, including LIN28B-TST in HCC and MARCO-TST in triple negative breast cancer, which exhibit therapeutic target potential^[22, 23]^. Furthermore, through systematic pan-cancer splicing junction profiling, we found that TSTs might generate neoantigens for immunotherapy^[22, 23]^. Notably, despite PDAC’s well documented splicing dysregulation^[24, 25]^, the systematic characterization of PDAC TSTs and their immunotherapeutic potential remains unexplored.

To overcome the limitations of neoantigen discovery in PDAC, we developed NeoAPP, a computational framework that systematically characterizes tumor-specific splicing junctions and exons to identify TSTs and their derived neoantigens. Applying NeoAPP across multiple PDAC cohorts, we uncovered that TST-derived neoantigens exhibit abundance and immunogenic potential compared to mutation-derived neoantigens, with distinct advantages in generating shared epitopes across patient cohorts. We also found that non-canonical splicing junction and transposable element produce high-yield neoantigen expansion, and alternative promoter usage is a key source of neoantigen-encoding TSTs (neoTSTs) regulated by FOXA2. Furthermore, we demonstrated potential packaging of neoTST into extracellular vesicles (EVs), positioning them as dual prognostic biomarkers and modulators of tumor-stromal crosstalk. Validation in HLA-A*02:01/HLA-A*11:01 transgenic mouse models confirmed antigen-specific CD8^+^ T cell responses, while therapeutic assessment in a syngeneic PDAC model revealed significant tumor growth inhibition mediated by neoTST vaccination.

## 2 Results

### 2.1 Computational framework of neoAPP for identification of TST-derived Neoantigens

To identify TST-derived neoantigens, we developed NeoAPP, which integrates three modules: specific splicing junction detection module, specific exon detection module and TST-derived neoantigen prediction module (**Figure 1A**). In NeoAPP, tumor RNA-seq data are systematically compared with reference splicing junction and exon profiles from normal tissues (GTEx, 3,178 samples across 29 tissue types by default). By applying stringent cutoffs and benchmarking each GTEx tissue independently, tumor-specific junctions and exons, and the resulting TSTs, are accurately identified (Figure S1A).

**Figure 1.**
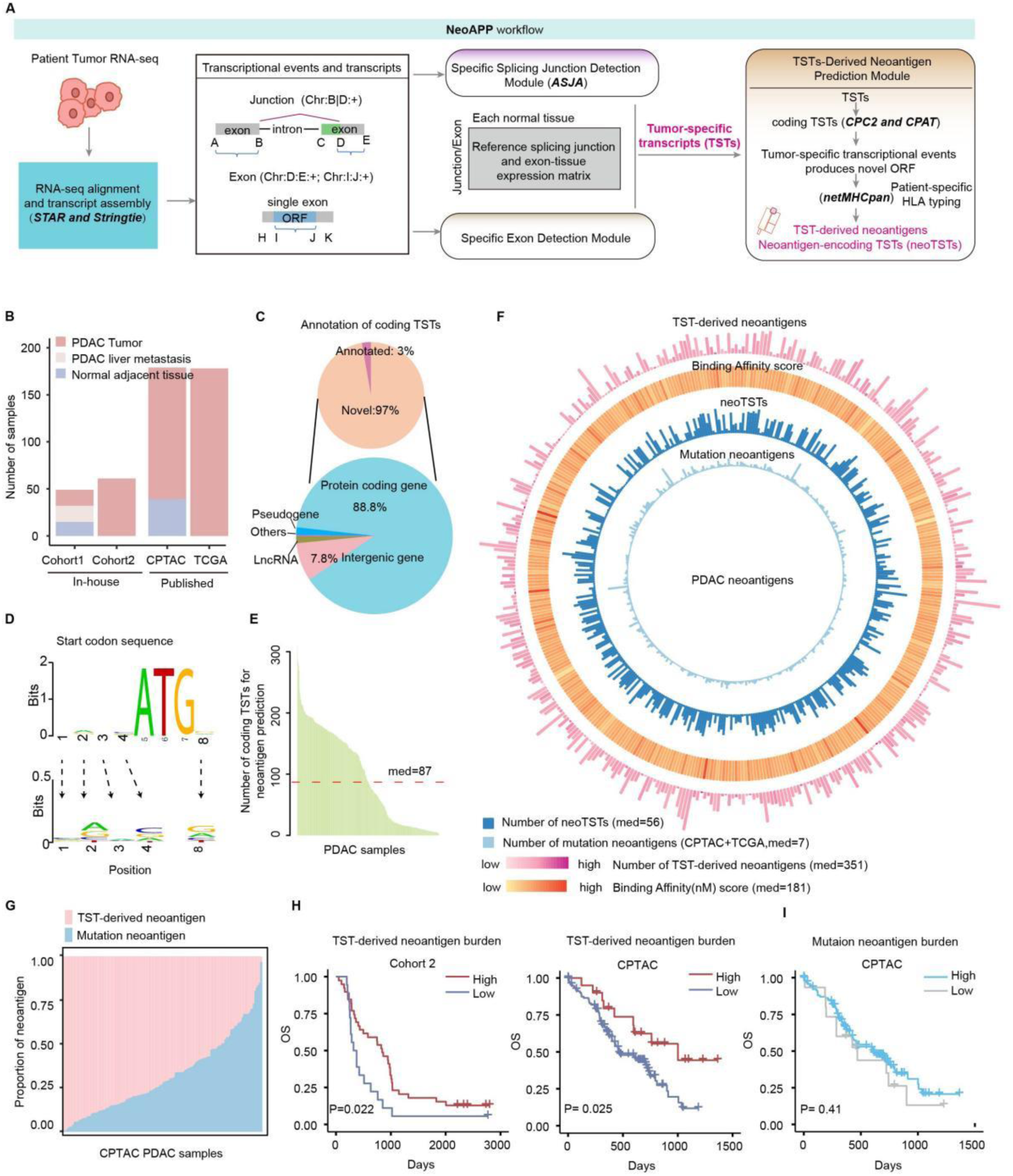
TST-derived neoantigen Landscape in PDAC Revealed by Multi-Cohort Transcriptomic Profiling. **(A)** NeoAPP workflow, including three main module: specific splicing junction detection module, specific exon detection module and TST-derived neoantigen preidction module. **(B)** Overview of the four PDAC RNA-seq cohorts. **(C)** Genomic annotation of coding TSTs. **(D)** Sequence motif analysis of translation start codons in coding TSTs (WebLogo visualization from the CPTAC cohort). **(E)** Bar plot showing the number of coding TSTs driven by tumor-specific transcriptional events (i.e., tumor-specific splicing junctions and exons). **(F)** Circos plot summarizing neoantigen features per PDAC sample: (1) number of TST-derived neoantigens; (2) binding affinity scores of TST-derived neoantigens; (3) number of neoantigen-encoding TSTs (neoTSTs); and (4) number of mutation-derived neoantigens. **(G)** Proportion of TST-derived versus mutation-derived neoantigens per sample in the CPTAC cohort. TST-derived proportion = TST-derived count / (TST-derived count + mutation-derived count); mutation-derived proportion = mutation-derived count / (mutation-derived count + TST-derived count). **(H)** Kaplan–Meier survival analysis stratified by TST-derived neoantigen burden (high vs. low) in Cohort 2 (left) and the CPTAC cohort (right). Log-rank test was used. The optimal cutoff point of TST-derived neoantigen burden was applied. **(I)** Kaplan–Meier survival curves for the CPTAC cohort stratified by mutation-derived neoantigen burden. The optimal cutoff point was used for stratification.

In the specific splicing junction detection module, we applied the ASJA algorithm to identify and quantify splicing junctions. Compared to approaches that rely on percent-spliced-in (PSI) values and group-based statistical comparisons, ASJA leverages transcript assembly to identify and quantify splicing junctions within individual samples and provide more unannotated junctions^[24]^. Unlike pooled-reference methods such as SNAF^[26]^, our tissue-stratified comparison against each GTEx tissue independently improves specificity in detecting tumor-specific events (Figure S1B). The exon detection module identifies tumor-specific exons by comparing genomic coordinates and expression patterns against the GTEx reference, thereby capturing unannotated single-exon transcripts. Finally, in the neoantigen prediction module, the coding potential of TSTs was assessed by in silico open reading frame (ORF) prediction. ORFs harboring tumor-specific junctions or exons were translated in silico, and peptides were evaluated for HLA class I binding affinity using NetMHCpan with patient-specific four-digit HLA typing. We define neoantigen-encoding TSTs (neoTSTs) as protein-coding TSTs that generate at least one predicted HLA-presented peptide. Accordingly, TST-derived neoantigens represent the complete set of candidate epitopes encoded by each neoTST.

#### TST-derived neoantigen landscape in PDAC revealed by multi-cohort transcriptomic profiling

To systematically characterize TST-derived neoantigen in PDAC, we analyzed 413 PDAC RNA-Seq samples across four cohorts (Figure 1C), including Cohort 1 (17 PDAC tumors with matched liver metastasis and 15 normal NAT samples, collected in-house), Cohort 2 (61 primary PDAC tumors, collected in-house), CPTAC cohort (140 PDAC tumors and 39 NAT samples), and TCGA cohort (178 PDAC tumors). Applying NeoApp (Figure S1C), we identified a median of 164 coding TSTs from 657 TSTs that derived from 317 tumor-specific splicing junctions and 488 tumor-specific exons per PDAC sample (Figure S1C). Genomic annotation revealed that 97% of coding TSTs were unannotated in GENCODE, primarily arising from alternative splicing of protein-coding genes (Figure 1C). Start codon motifs in these transcripts exhibited Kozak sequence compatibility (Figure 1D), suggesting stable translational potential. To prioritize high-confidence candidates, we retained only open reading frames directly supported by tumor-specific splicing junctions or exons for neoantigen prediction, yielding a median of 87 coding TSTs per sample (Figure 1E). Finally, from 401 PDAC samples with HLA typing data, we identified a median of 351 neoantigens, with a median MHC I binding affinity of 181 nM, derived from a median of 56 neoTSTs per sample (Figure 1F).

We also analyzed somatic mutation-derived neoantigens using whole-exome sequencing (WES) data from the CPTAC cohort and curated mutation neoantigens from the TSNAdb database^[27]^ for TCGA cohort. This analysis revealed a median of 7 mutation-derived neoantigens per tumor (range: 0-51, excluding 3 hypermutated cases with non-synonymous somatic mutations >300), derived from 27 median non-synonymous somatic mutations per sample. Notably, TST-derived neoantigens exhibited higher abundance than mutation neoantigens (Figure 1F), with per-sample ratios of the two classes demonstrating mutually exclusive dominance, thereby compensating for immunogenic epitope scarcity (Figure 1G, Figure S1D). Critically, elevated TST-derived neoantigen burden correlated with improved overall survival (OS) in both in-house (Cohort 2) and external validation cohorts (CPTAC/TCGA; log-rank p < 0.01; Figure 1H, Figure S1E), whereas mutation-derived neoantigens showed no prognostic significance (Figure 1I, Figure S1E).

### 2.2 Proteomic validation and immunogenic primacy of TST-derived neoantigens in PDAC

To validate the translational potential of neoTSTs, we performed integrated proteogenomic analysis of matched RNA-seq and mass spectrometry (MS) data. Across 140 PDAC tumors, 17.3% of neoTSTs per patient demonstrated peptide-level evidence (**Figure 2A**), with representative PDIA3-derived neoTSTs showing concordant exon-skipping patterns in transcriptomic and MS profiles (Figure 2B-C). TST-derived neoantigens exhibited enhanced immunogenic properties compared to mutation-derived counterparts: 15.8% demonstrated promiscuous HLA-I binding to ≥ 2 alleles (4-digit resolution) (Figure 2D, Figure S2A) with enhanced median affinity (IC50 = 141.02 nM; Figure S2B), and recurrence rates (1.5% vs 0.5%) and patient coverage increased 3-fold (Figure 2E). For instance, the TST-derived neoantigen QANSFPLTF (chr2:89040462|89046412) targeted 105 patients versus 21 for the KRAS G12V-derived epitope (Figure 2F). The top 10 TST-derived neoantigens collectively engaged 41.4% (171/413) of patients, each binding 20 HLA alleles on average (IC50 range: 8.57-494.47 nM; Figure 3G), underscoring their potential as universal immunotherapeutic targets.

**Figure 2.**
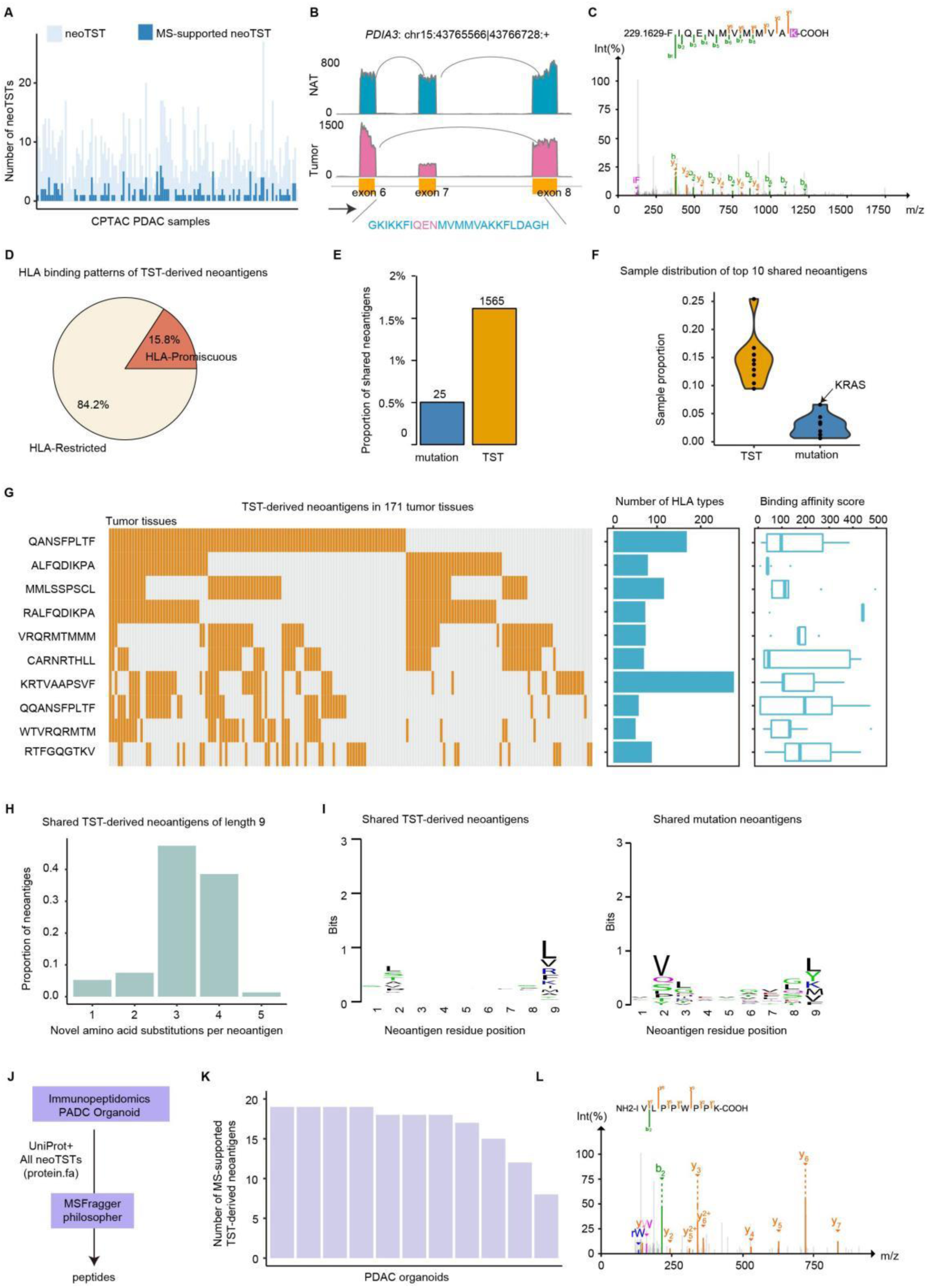
Proteomic validation and immunogenic primacy of TST-derived neoantigens in PDAC. **(A)** Number of neoTSTs identified in CPTAC PDAC samples by integrating transcriptomic and proteomic data. **(B)** Sashimi plot showing an exon-skipping splicing junction in PDIA3 (chr15:43765566|43766728). **(C)** Mass spectrometry (MS) spectrum of the FIQENMVMMVA peptide. **(D)** Pie chart showing the proportion of TST-derived neoantigens binding to promiscuous HLA alleles (i.e., ≥2 HLA alleles) versus those binding to allele-specific HLAs. **(E)** Number of shared neoantigens across samples or cohorts (TST-derived: present in ≥2 cohorts; mutation-derived: present in ≥2 samples). **(F)** Proportion of samples harboring each of the top 10 shared neoantigens. **(G)** Tissue distribution of the top 10 shared TST-derived neoantigens (waterfall plot, left), corresponding HLA binding diversity (middle), and predicted binding affinity (right). **(H)** Proportion of novel amino acids (not annotated in UniProt) in 9-mer shared TST-derived neoantigens. **(I)** Sequence motifs of shared TST-derived versus mutation-derived neoantigens (WebLogo visualization). **(G)** Workflow for immunopeptidomics analysis. **(K)** Sashimi plot showing generation of the MEDFRKNFL peptide from a single exon event. **(L)** MS spectrum of the MEDFRKNFL peptide.

**Figure 3.**
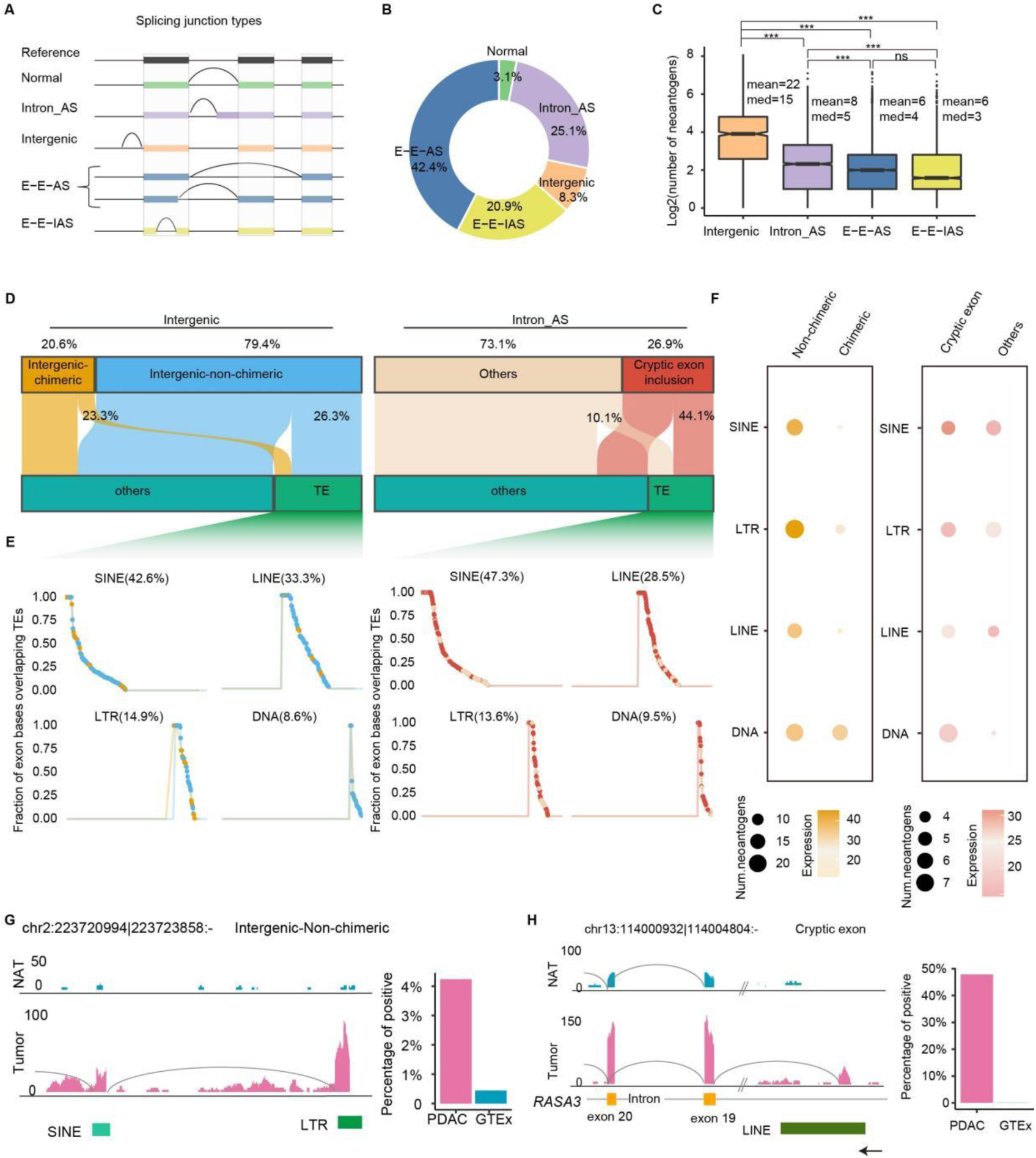
Non-canonical splicing and transposable element generate high-yield neoantigen expansion in PDAC. **(A-B)** Classification and proportions of splicing junction types. **(C)** The number of TST-derived neoantigens derived by different splicing junction types. Wilcoxon rank-sum test, *P<0.05, **P<0.01, ***P<0.001. **(D)** Proportion of exons generated from tumor-specific splicing in intergenic and intronic regions overlapping with TE (Sankey diagram). **(E)** Proportion of bases derived from TE sequences in exons generated by tumor-specific splicing. Each dot represents one aberrant splicing event, where blue indicates intergenic non-chimeric splicing junctions, orange indicates intergenic chimeric splicing junctions, dark red indicates intronic cryptic exon inclusion events, and light pink indicates other intronic splicing junctions. **(F)** Point plot showing the average neoantigen derived from different neoTST subtypes. **(G-H)** Representative Sashimi plots of intergenic-non-chimeric and cryptic exon inclusion. Right: expression frequency of tumor-specifc junction driving neoTST.

TST-derived neoantigens harbored more novel amino acid residues per epitope than mutation-derived counterparts (median = 3 vs 1; Figure 2H), a feature associated with enhanced immunogenicity due to greater divergence from self-peptides and reduced susceptibility to central tolerance mechanisms. The motif analysis of all TST-derived shared neoantigens revealed their sequence characteristics, including absence of valine bias at position 2 and C-terminal leucine enrichment (Figure 2I), suggesting differences in the strength and type of MHC molecules bound to the mutation neoantigens. Immunopeptidome profiling of 10 PDAC organoids, performed by re-analyzing public raw data from the MSV000096853 dataset^[27]^, confirmed the presentation of TST-derived neoantigens, with a median of 16 computationally predicted candidates detected per sample (Figure 2J-K). For example, the 9-mer peptide IVLPPWPPK, derived from a neoTST “chr18:51067187 – 51101104” (Figure 2L). Collectively, TST-derived neoantigens feature conserved expression across patients, broad HLA presentation, and unique sequence features, positioning them as a potential target reservoir for PDAC immunotherapy.

### 2.3 Non-canonical splicing and transposable element generate high-yield neoantigen expansion in PDAC

Our analysis demonstrated that multi-exonic neoTSTs accounted for the vast majority (96.6%) of neoantigens, whereas single-exon neoTSTs represented only a minor fraction (3.3%) (Figure S3A). To elucidate the diversity of multi-exonic neoTSTs, we classified splicing junctions into five distinct categories: canonical splicing (normal, 13.3%), intron alternative splicing (intron_AS; 25.1%), intergenic splicing (8.3%), exon-exon alternative splicing (E-E-AS; 42.4%), and exitron splicing (E-E-IAS; 20.9%) (**Figure 3A-B**). Representative examples of E-E-AS and E-E-IAS types are illustrated in Figure S3B-C. Transcripts derived from intergenic and intron_AS regions exhibited paradoxical properties: their ORFs averaged 535 and 1,066 nucleotides, respectively-significantly shorter than UniProt-annotated proteins (median 1,542 nucleotides; Figure S3D). Nevertheless, these truncated ORFs demonstrated exceptional neoantigen productivity, with intergenic and intron_AS-derived neoTSTs generated averages of 22 and 8 neoantigens respectively, significantly higher yields compared to other spliced variants (6 neoantigens, P<0.001, Figure 3C).

Building on the established role of transposable elements (TEs) in generating aberrant splicing isoforms, we systematically mapped their contributions to neoTST across intergenic and intronic regions. Intergenic splicing events were stratified into chimeric (20.6%) and non-chimeric isoforms (79.4%), while intronic alternative splicing (intron_AS) events were categorized into cryptic exon inclusion (26.9%) and other anomalies (71.3%) (Figure 3D, Figure S3E-F). Notably, canonical GT-AG splice sites drove cryptic exon incorporation in intron_AS events (e.g., chr12:26922700|26924051; Figure S3G-H), which were functionally enriched in pathways such as protein digestion and absorption (Figure S3I). TE overlap analysis revealed positional specificity: 26.3% of non-chimeric intergenic neoTSTs and 23.3% of chimeric isoforms harbored TE-derived sequences, with intronic cryptic exons exhibiting the highest TE association (44.1%; Figure 3D). Moreover, SINE elements dominated TE contributions in both intergenic (42.6%) and intronic (47.3%) regions, followed by LINE and LTR elements, highlighting subtype-specific retrotransposon activity (Figure 3E). On average, exons overlapping with TEs contained 40% TE-derived nucleotides, underscoring the substantial contribution of retrotransposon insertions to the expansion of the neoantigenic repertoire. Quantitative analyses further revealed that TE-associated neoTSTs yielded significantly higher neoantigen output per transcript compared to their non-TE counterparts. Specifically, LTR-associated intergenic non-chimeric neoTSTs generated an average of 22 neoantigens per transcript, while SINE-associated cryptic exons produced approximately 5 neoantigens per neoTST (Figure 3F). Recurrent examples included the LTR-driven intergenic neoTST chr2:223720994|223723858 detected in 4% tumor samples (Figure 3G). High-prevalence neoTSTs such as LINE-driven neoTST chr13:114000932|114004804 (47.8%) exhibited broad patient-level recurrence (Figure 3H). Furthermore, single-exon neoTSTs with nearly 40% TE overlap expanded the antigenic repertoire, with DNA-associated isoforms exhibiting 1.5-fold higher neoantigen yields than non-TE counterparts (Figure S3J-K). These data collectively indicate that TE-associated neoTSTs substantially expand the neoantigen landscape, offering a novel genomic reservoir for immunotherapeutic targeting.

### 2.4 Alternative promoter usage as a key source of neoTSTs regulated by FOXA2

Alternative promoter usage through transcriptional start site (TSS) selection represents a critical mechanism driving transcriptome diversity in cancer^[30]^. To investigate whether alternative TSS activation contributes to the emergence of neoTSTs in PDAC, we systematically mapped splicing junction distributions relative to TSS positions. This analysis identified 2,061 first-junction-derived neoTSTs (F-neoTSTs) as dominant contributors to neoantigen production (**Figure 4A**). These transcripts predominantly originated from two genomic contexts: intergenic splicing events (n=604) and intron_AS events (n=549) (Figure 4B). F-neoTSTs exhibited exceptional immunogenic potential, producing an average of 13 neoantigens per transcript over other splicing junction-derived neoTSTs (Figure S4A). The TSSs of F-neoTSTs were predominantly marked by H3K27ac and H3K4me3 within ±1 kb regions (Figure 4C), with 69% arising from previously unannotated (Figure 4D), indicative of active alternative promoter usage. These findings establish that novel TSS selection expands the neoTST repertoire.

**Figure 4.**
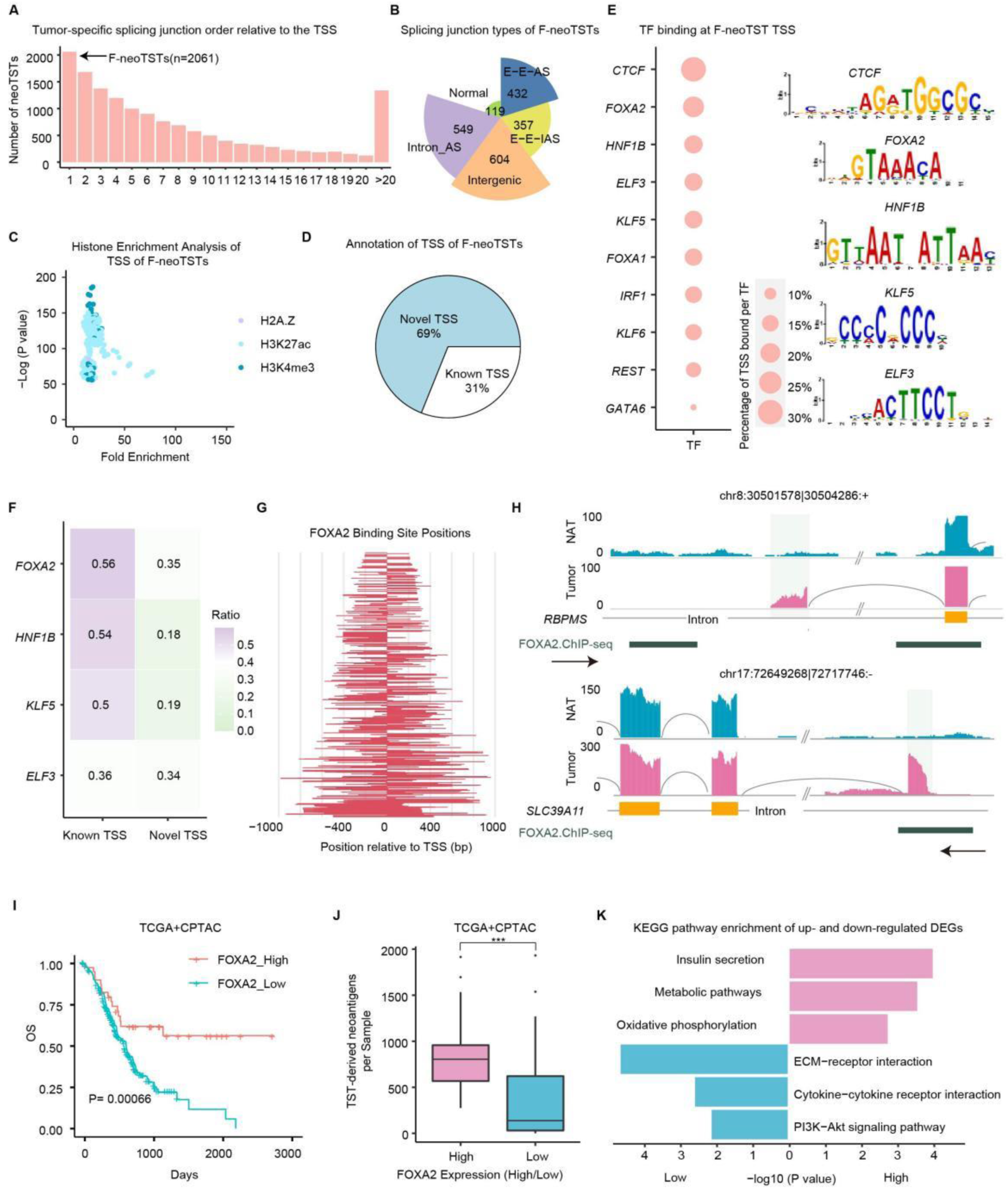
Alternative promoter usage as a Key Source of neoTSTs Regulated by FOXA2. **(A)** Spatial distribution of tumor-specific splicing junctions driving neoTSTs relative to the TSS. **(B)** Pie chart showing the distribution of splicing junction types that generate F-neoTSTs. **(C)** Point plot displaying TSS histone enrichment for F-neoTSTs, analyzed using ChIP-Atlas. **(D)** Pie chart comparing the novel and known TSS of F-neoTSTs. **(E)** Ranking of TFs by binding sites frequency at F-neoTST TSSs, alongside motif analysis of top TFs using SEA enrichment (right). **(F)** Proportion of known and novel TSSs bound by TFs, with normalization performed separately for known and novel TSS subsets. **(G)** Linear plot mapping FOXA2 binding sites relative to TSS. **(H)** Sashimi plot visualizing examples of F-neoTSTs regulated by FOXA2. “FOXA2-ChIP-seq” denotes ChIP-seq of FOXA2 indicating the binding regions in pancreatic cancer cell lines. **(I)** Kaplan-Meier survival analysis (log-rank test) comparing overall survival of PDAC samples (TCGA and CPTAC cohorts) stratified by optimal cutoff point of FOXA2 expression levels. **(J)** Box plot showing differences in TST-derived neoantigen counts between samples with high and low FOXA2 expression.Wilcoxon rank-sum test, *P<0.05, **P<0.01, ***P<0.001. **(K)** KEGG enrichment analysis of differentially expressed genes. Blue represents the FOXA2 low group, and pink represents the high group.

Integrative ChIP-seq analysis revealed binding sites of transcription factors (TFs) at TSS±1 kb of F-neoTSTs, including CTCF, FOXA2, HNF1B, KLF5, and ELF3 (Figure 4E). MEME SEA motif analysis further confirmed the presence of these TF binding motifs in the corresponding TSS regions (Figure 4E). Among them, FOXA2 emerged as the master transcriptional regulator, binding to 452 F-neoTST promoters (Figure 4E) and preferentially regulating novel TSSs (35%; Figure 4F). FOXA2 binding localized within ± 400 bp of core promoter regions (Figure 4G), exemplified by two intron_AS-derived F-neoTSTs: chr8:30501578|30504286 and chr17:72649268|72717746 (Figure 4H). Clinically, elevated FOXA2 expression correlated with improved overall survival (log-rank P < 0.01; Figure 4I, Figure S4B), consistent with its tumor-suppressive role in PDAC^[31]^. FOXA2-high tumors exhibited elevated TST-dervied neoantigen levels (Figure 4J), suggesting the potential role in immunogenic antigen biogenesis. Pathway analysis further linked FOXA2 to metabolic remodeling, including activation of insulin secretion and suppression of PI3K-Akt and cytokine-cytokine receptor signaling (Figure 4K, Figure S4C). Collectively, these findings suggest F-neoTSTs as pivotal contributors to the PDAC neoantigen landscape, with FOXA2 orchestrating both their transcriptional activation and anti-tumor immune priming.

### 2.5 Distribution of neoTSTs in tumor microenvironment and extracellular vesicles

We further analyzed single-cell RNA-seq data of PDAC to explore the expression of these neoTSTs in tumor tissues. Dimensionality reduction and clustering revealed distinct cellular populations, including tumor epithelial cells, fibroblasts, stellate cells, schwann cells, endothelial cells, mast cells, neutrophils, macrophages, B cells, and T cells (Figure S5A-B). Using SCASL, a computational tool for splicing junction detection, we identified cells (gene counts > 2,000) expressing single-cell neoTST (sc-neoTST) (**Figure 5A**). After stringent filtering, 19.2% of tumor cells harbored sc-neoTST (Figure 5B) to disclose neoTSTs were tumor specific in single cell level. Interestingly, single-cell analysis demonstrated that 4.7% of fibroblasts expressed sc-neoTSTs, representing the highest frequency across all non-malignant cell types analyzed (Figure 5C, Figure S5C). Given the role of extracellular vesicles (EVs) in cell-cell communication^[32]^, we hypothesize that EVs might transfer neoTSTs to fibroblast cells. Supporting with this notion, many EVs related pathways were higher expression in expressed neoTSTs in tumor cell than non-expressing cells, especially the endosomal sorting complex required for transport (ESCRT) pathway (Figure 5D), a key driver of EVs biogenesis. Conversely, fibroblasts harboring sc-neoTSTs exhibited upregulated pathways associated with EVs uptake (e.g., Pinocytosis) and Vesicle Docking (Figure 5E), suggesting enhanced EVs internalization capacity. Further analysis of fibroblast subtypes signature expression suggested myCAF-like subsets (e.g., MMP1+Fib, LRRC15+Fib) as the predominant recipients of EV cargo (Figure 5F, Figure S5D).

**Figure 5.**
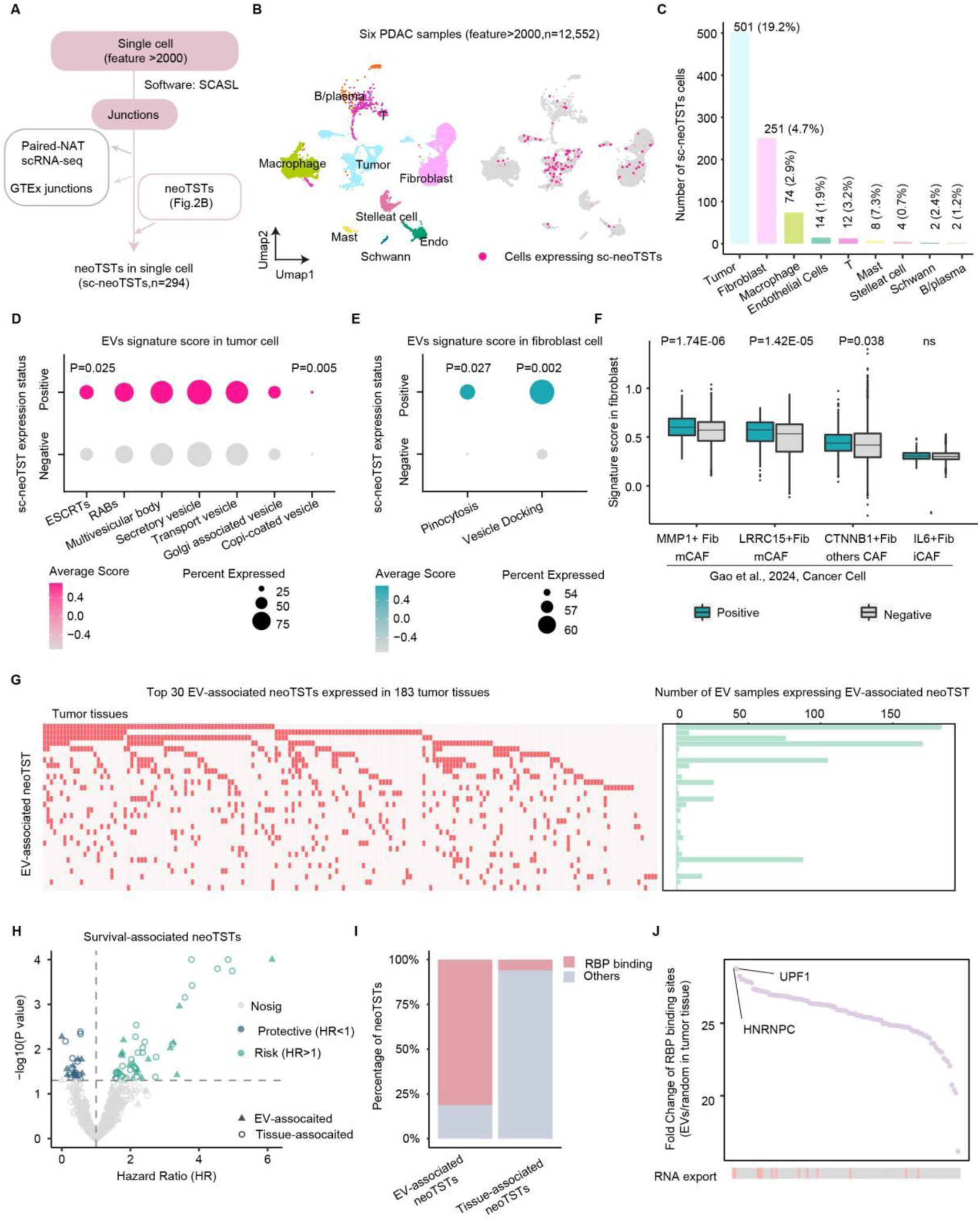
Distribution of neoTSTs in tumor microenvironment and extracellular vesicles. **(A)** Flowchart outlining the neoTSTs detection workflow in single-cell RNA-seq. **(B)** UMAP projection of PDAC tumor single-cell data, colored by cell type (left) and UMAP projection of PDAC single-cell data, with cells expressing sc-neoTSTs highlighted in deep pink and negative cells shown in grey (right). **(C)** Bar plot displaying number of single cell expression of sc-neoTSTs. **(D)** Dot plot evaluating EVs signature scores in tumor cells with versus without sc-neoTST expression. **(E)** Dot plot evaluating EVs signature scores in fibroblast cells with versus without sc-neoTST expression. **(F)** Box plot assessing fibroblast signature scores in fibroblast cells harboring sc-neoTST compared to negative counterparts. **(G)** Waterfall plot ranking the top 30 neoTSTs detected in EVs by their tissue prevalence (left), alongside a bar plot quantifying their detection frequency across EV samples (right). **(H)** The Volcano plot showing the survival-associated neoTST. **(I)** Proportional bar chart illustrating RBP binding site enrichment on neoTSTs in EVs compared to tissue sample. **(J)** Ranking of RBPs by the fold change in binding site counts for neoTSTs between EVs and tissue. The red line indicates that RBPs are associated with RNA export.

To further evaluate whether EVs carry neoTSTs, we analyzed matched EVs and tumor tissue RNA-seq data from 57 PDAC patients (Cohort 2). Strikingly, 61.4% of cases (n=35) exhibited concordant detection of neoTSTs in paired EVs (Figure S5E), confirming their active packaging into EVs. Expanding to a large plasma EV RNA-seq cohort of PDAC (n=373), we identified 1,705 EV-associated splicing junctions linked to neoTSTs (Figure S5F). The top 30 EV-associated neoTSTs (ranked by expression frequency in tumor samples) exhibited high prevalence across both tumor tissues and extracellular vesicles (Figure 5G). In addition, survival analysis identified 51 risk-associated and 23 protective neoTSTs with overall survival (log-rank P < 0.05), 31.8% of which were detectable in EVs (Figure 5H). For example, the high-risk neoTST chr3:185524160|185533571 (detected in 141 PDAC samples) correlated with shortened survival (HR=1.80, P=0.0063), while the protective splicing junction chr17:3935268|3936267 (detected in 40 samples) was associated with improved outcomes (HR=0.47, P=0.017). These findings suggest that, beyond their immunogenic potential, neoTSTs may also reflect underlying tumor biology or disease state, thereby contributing to their association with patient survival as potential biomarkers. To dissect the selective incorporation of neoTSTs into EVs, we investigated RNA-binding protein (RBP)-mediated regulation by integrating ENCODE eCLIP-seq data. Approximately 80% of neoTSTs in EVs harbored RBP binding sites-significantly enriched compared to tissue-associated neoTSTs (Figure 5I). By comparing the number of binding sites between EV-associated and tissue-associated neoTSTs, we identified HNRNPC and UPF1 as dominant mediators of neoTST loading into EVs (Figure 5J), implicating RBP regulation in EV cargo selection.

### 2.6 neoTST elicited antigen-specific CD8+ T cell responses in HLA transgenic mice and inhibited tumor growth in a syngeneic PDAC model

To validate the endogenous immunogenicity, we systematically screened 10 highly expressed and cross-sample-conserved neoTSTs in PDAC (**Figure 6A-B**, Figure S6A). NetMHCpan predictions identified at least one HLA-A*11:01-restricted epitope across all candidates, with six neoTSTs additionally harboring HLA-A*02:01-binding epitopes (Figure S6B). Accordingly, HLA-A*11:01 and HLA-A*02:01 humanized transgenic mice received two intramuscular doses of a mixed neoTST mRNA-LNP vaccine (5 μg/dose). One week post-immunization, splenocytes were transfected with neoTST mRNA and subjected to IFN-γ ELISpot and flow cytometry to quantify antigen-specific T cell responses (Figure 6C). Results demonstrated that 40% of neoTSTs elicited robust HLA-A*11:01-restricted T cell activation, while 50% triggered HLA-A*02:01-dependent responses (Figure 6D-G, Figure S6C-D). Intriguingly, three neoTSTs activated dual HLA-restricted T cells, indicating multi-HLA presentation capacity.

**Figure 6.**
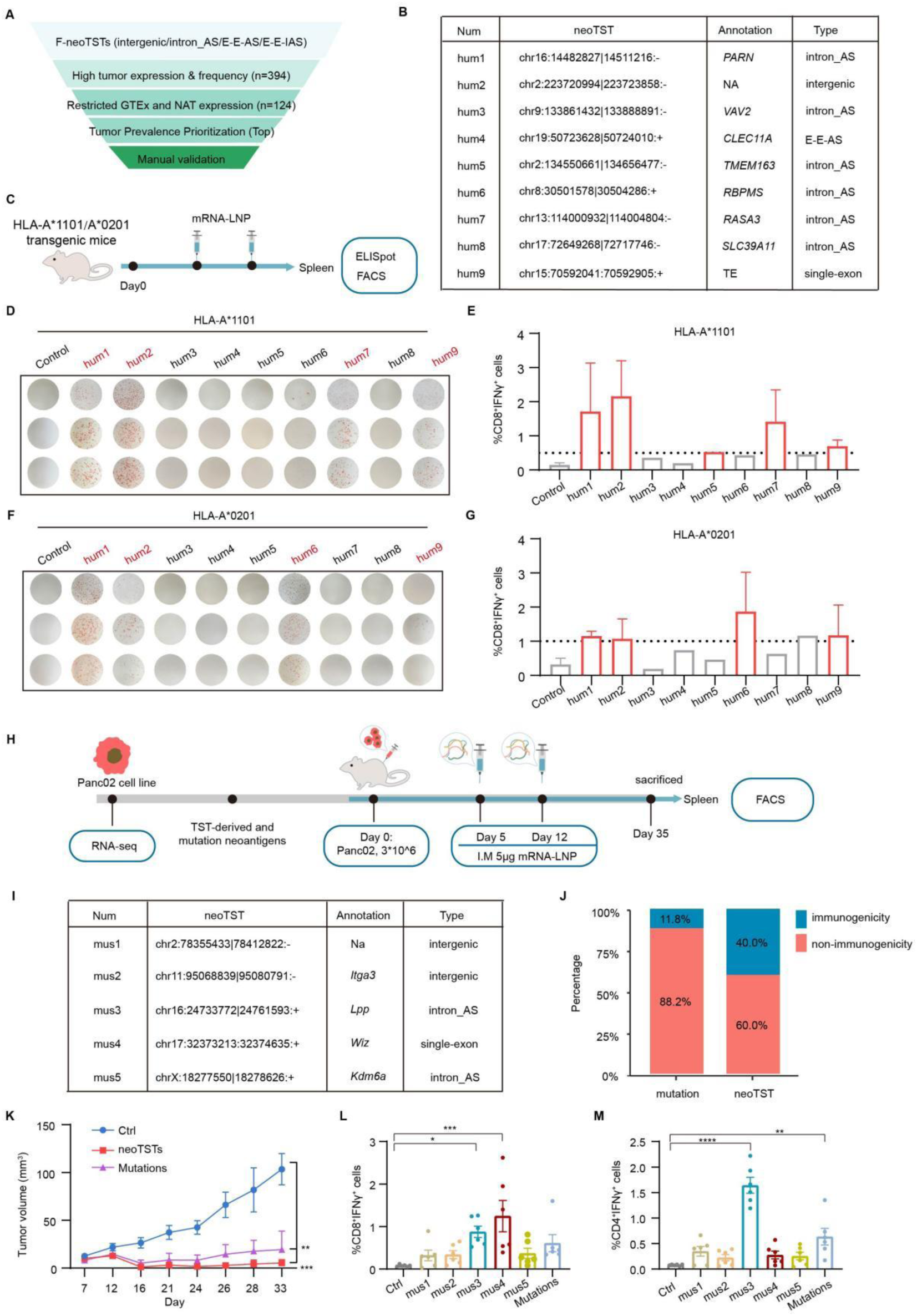
neoTST elicited antigen-specific CD8+ T cell responses in HLA transgenic mice and inhibited tumor growth in a syngeneic PDAC model. **(A)** Screening of representative PDAC neoTSTs. **(B)** Characterization of representative PDAC neoTSTs. **C,** Workflow for immunogenicity validation in humanized HLA transgenic mice. **(D-G),** HLA-A*11:01 and HLA-A*02:01 transgenic mice were immunized with mixed neoTST mRNA-LNP(C). One week post-immunization, splenocytes were harvested and incubated with individual neoTST mRNA-LNP ex vivo and monitored by ELISpot assays. Empty mRNA-LNP served as negative control, respectively. ELISpot images (D,F) and reactive CD8⁺ T cell populations isolated by FACS (E,G) are shown. **(H)** Schematic of using the murine pancreatic cancer cell line Panc02 as a model system to validate *in vivo* immunogenicity of neoTSTs and assess their therapeutic potential (n = 6). **(I)** Characterization of representative neoTSTs in Panc02. **(J)** Proportion of immunogenic neoantigens: Mutation-derived versus neoTST. **(K)** Tumor growth kinetics analyzed by one-way ANOVA. Data are shown as mean± SEM. *P<0.05, **P<0.01, ***P<0.001. Control: unvaccinated mice. n=6. **(L-M)** Proportions of CD8+ T cells (l) and CD4+ T cells (m) induced by neoTST versus mutation-derived neoantigens. One-way ANOVA was used; data shown as mean±SEM. *P<0.05, **P<0.01, ***P<0.001. Control: unvaccinated mice. n=6.

To validate the *in vivo* immunogenicity of neoTSTs and assess their therapeutic potential, we utilized the murine pancreatic cancer cell line Panc02 as a model system (Figure 6H). Five representative neoTSTs were selected for testing in the Panc02 murine pancreatic cancer model (Figure 6I). To evaluate therapeutic efficacy, mice bearing subcutaneous Panc02 tumors were vaccinated with a pool of five synthesized neoantigen sequences on days 5 and 12. For comparison, we also identified mutation-derived neoantigens in Panc02 cells. We detected 17 nonsynonymous single nucleotide substitutions that resulted in novel protein sequences, which were considered potential neoantigens based on rigorous criteria (Figure S6E). The mRNAs of corresponding mutational sequence (25 amino acids each) were synthesized for immunogenicity testing in mice. The mutation-derived neoantigens were administered as an experimental group under identical dosing and timing as the neoTST group. In stark contrast, while merely 11.8% (2/17) of mutational neoantigens activated detectable CD8+ T cell populations, TST-derived neoantigens demonstrated significantly higher immunogenicity with a 40% (2/5) response rate (Figure 6J, Figure S6F-G).

Experimental analyses revealed significant tumor suppression in both neoTST and mutation neoantigen vaccine groups compared to controls by day 35 (Figure 6K, Figure S6H). Notably, neoTST-based vaccines exhibited a favorable trend in tumor regression efficacy than mutation neoantigen formulations. Flow cytometric profiling of splenocytes showed that neoTSTs induced antigen-specific CD8+ T cell populations, with a particularly strong response observed against the mus4 neoTST (a novel single-exon) (Figure 6L). Both the mu3 neoTST (a cryptic exon) and mutation-derived neoantigens significantly activated antigen-specific CD4+ T cells (Figure 6M). Our data provide experimental evidence that neoTSTs can impede tumor growth and induce specific T cell responses, underscoring their potential as a novel immunotherapeutic strategy for PDAC.

## 3 Discussion

Recent clinical trials have demonstrated the therapeutic promise of mutation-derived neoantigen vaccines in resected PDAC. However, the scarcity of actionable mutations in this genomically stable malignancy highlights the need for alternative antigen sources. To address this gap, we developed NeoAPP, a dedicated computational framework designed to systematically detect TSTs by integrating exon- and junction-level analyses across individual tumor samples. NeoAPP enables sensitive and reliable identification of TSTs and their derived neoantigens, offering a tool to uncover novel antigenic targets beyond mutation-based sources. Applying NeoAPP, we systematically identify TST-derived neoantigens as a potent yet underexplored reservoir of immunogenic targets in PDAC. Our results showed that a median of 351 neoantigens from 56 neoTSTs per sample. These originate from diverse genomic events, including TE-associated intronic exons, and intergenic splicing. Alternative promoter usage, primarily regulated by FOXA2, plays a key role in neoTST generation. Notably, bioinformatic analysis showed that neoTSTs are detectable in EVs and transferred to the tumor microenvironment, particularly to myCAFs. Functional assays in HLA-A transgenic models confirmed neoTST-induced CD8⁺ T cell responses, while Panc02 studies demonstrated reduced tumor growth and activation of both CD8⁺ and CD4⁺ T cells. We uncover a vast repertoire of therapeutically actionable TST-derived neoantigens, advancing novel immunotherapeutic strategies for this recalcitrant cancer. Our identification of neoantigens derived from tumor-specific splicing events aligns with recent breakthroughs in immunopeptidomics, which revealed cryptic antigens as potent targets for T cell recognition in pancreatic cancer^[28]^. While Ely et al. emphasized non-canonical peptides from translational dysregulation, our study highlights transcriptional aberrations, generate a complementary reservoir of shared and immunogenic neoantigens. Both approaches highlight the limitations of relying solely on somatic mutations for neoantigen discovery in PDAC and underscore the therapeutic potential of targeting non-canonical antigen sources. The convergence of these findings advocates for integrated multi-omic strategies to exploit the full spectrum of tumor-specific antigens in genomically stable cancers.

The concept that aberrant mRNAs can generate neoantigens has gained increasing attention. Computational tools such as SPLICE-neo^[33]^, SNAF^[26]^, and IRIS^[34]^, have been developed to identify splicing-derived neoantigens. However, most rely solely on junction-level analysis, limiting their ability to comprehensively capture tumor-specific transcript isoforms. In contrast, NeoAPP integrates splicing junctions and exon-level genomic positions with expression data, enhancing transcript validation and enabling the identification of single-exon TSTs often missed by junction-only approaches. For example, in a murine PDAC model, we identified a novel single-exon transcript that elicited strong CD8⁺ T cell responses, underscoring its immunogenicity. Besides, ASJA tool using in specific splicing junction detection module, extracts junctions from reference-guided transcript assemblies, ensuring higher reliability. Unlike conventional splicing tools that focus on isoforms derived from a GTF annotation^[35]^ and require group comparisons^[36]^, ASJA enables single-sample detection, quantification, and cross-sample comparison of all junctions. This strategy captures a large fraction of unannotated splice sites, providing a more comprehensive view of transcriptome complexity. Additionally, unlike tools that apply global fold-change (e.g., SNAF) or expression thresholds (e.g., IRIS), our single-tissue framework benchmarks against 29 GTEx tissues and paired normal samples, improving specificity by controlling for normal tissue transcriptional baselines and helps avoid interference from genes with tissue-specific expression in normal.

The development of universal neoantigens offers clinical advantages over personalized vaccines, including reduced cost and faster production. While the canonical KRAS neoantigen VVGAVGVGK was predicted to bind MHC I in only 6.9% of patients (21/301), our TST-derived neoantigens showed broader predicted population coverage. We identified 1,565 potentially shared neoantigens (1.5% of total), including QANSFPLTF with MHC I binding in 25.4% of cases (105/413). The top 10 neoantigens collectively covered 41.4% of patients, highlighting their promise in low-TMB tumors like PDAC where shared mutation-derived epitopes are rare. These findings parallel glioma studies in which conserved splicing-derived antigens such as GNAS neojunctions exhibit broad immunogenicity. Moreover, PDAC-derived TST neoantigens were more abundant and enriched in novel amino acid sequences—a feature associated with increased immunogenicity ^[37]^. These parallels underscore transcriptional dysregulation as a pan-cancer mechanism for generating immunogenic epitopes, particularly in low-TMB malignancies like PDAC.

While TSS plasticity is known to drive isoform diversity in cancer^[38, 39]^, our study reveals its overlooked role in shaping the tumor immunopeptidome. We found that 69% of first-junction-derived F-neoTSTs arise from novel TSSs, marked by epigenetically active chromatin states. These unannotated TSSs generate novel ORFs with neoantigen potential. Notably, FOXA2—a transcription factor associated with favorable survival—binds near 35% of these novel TSSs. FOXA2 not only suppresses oncogenic PI3K signaling but also facilitates cryptic TSS activation, thereby promoting F-neoTST generation. Although mechanistic studies are warranted, our multi-omic data demonstrate the central role of FOXA2 and TSS regulation in expanding the neoantigenic landscape.

Beyond cell-autonomous antigen presentation, our analyses implicate EVs in disseminating neoTSTs to myofibroblast-like CAFs, a subset of stromal cells critical for PDAC progression^[40]^. This observation of EVs is increasingly recognized as key players in shaping the tumor microenvironment through the transfer of bioactive molecules in pancreatic cancer^[41]^. While the fate of EV-transferred neoTSTs—whether translated or processed for MHC presentation—remains unclear, their enrichment in EVs suggests a mechanism for systemic immune priming or stromal reprogramming. For instance, You et al. demonstrated that aberrantly expressed repetitive RNAs in PDAC are transferred to cancer-associated fibroblasts via EVs, triggering type I interferon responses in CAFs and subsequently reprogramming their phenotypic states^[42]^. Future studies employing spatial transcriptomics and co-culture models could elucidate whether EV-mediated neo-TST transfer enhances broad HLA presentation or induces tolerance.

Although our analytical approach identified a large repertoire of TST-derived neoantigens, providing reliable targets for mRNA-based therapies in pancreatic cancer, several limitations of our study should be acknowledged. First, current limitations in HLA-II binding prediction tools prevented the assessment of MHC class II-restricted neoantigens in our study. Additionally, while our murine model provides robust proof-of-concept for the immunogenicity of TST-derived neoantigens, the translatability of these findings to human PDAC requires validation in clinical cohorts. Future studies should focus on validating these findings in patient-derived models and exploring the potential of TST-derived neoantigens in clinical immunotherapy.

## 4 Experimental Section PDAC multi-cohort source

We analyzed four independent PDAC cohorts to identify TST-derived neoantigens. Cohort 1 (n = 34 tumors; n = 15 NAT samples) and Cohort 2 (n = 61 tumors) were prospectively collected in-house. Formalin-fixed, paraffin-embedded samples from Cohorts 1 and 2 were obtained and sequenced as described in the previous study^[42]^. Publicly available RNA-seq BAM files for CPTAC-PDAC (n = 140 tumors; n = 39 NAT samples) and TCGA-PAAD (n = 178 tumors) were downloaded from the NCI Genomic Data Commons (GDC) using the gdc-client tool under controlled-access authorization. In addition, RNA-seq BAM files from 29 normal tissue types (GTEx v8) were obtained via dbGaP and used as control datasets.

### RNA-seq Processing

For Cohorts 1 and 2 with raw sequencing data, quality control was performed using FastQC, followed by alignment to the GRCh38.p12 reference genome using STAR^[44]^ (v2.5.3a) in two-pass mode with chimeric junction detection enabled. Transcript assembly for each sample was conducted using StringTie^[45]^ (v2.2.1) with guidance from GENCODE v29 annotations. For the CPTAC and TCGA cohorts, where aligned BAM files were available, transcript assembly was performed directly using StringTie (v2.2.1) in reference-guided mode. The same processing pipeline was applied to BAM file of GTEx normal tissues and NAT samples to ensure consistency in transcriptome annotation across all cohorts.

### Reference junction and exon-tissue expression matrix

To identify tumor-specific transcripts (TSTs) that defined as transcripts generated from tumor-specific splicing junctions or exons, we first quantified splicing junctions and exons across 29 GTEx normal tissue types to create a reference profiles. Splicing junctions were identified using ASJA^[46]^, and expression was quantified as coverage per ten million reads (CPT). Exons were annotated and quantified from StringTie-assembled transcripts, and expression was normalized as e_ncov.

Next, we established reference expression matrices for both splicing junctions and exons using 29 GTEx normal tissue. For each tissue type, we calculated three parameters for every splicing junction and exon: (a) median expression value, (b) frequency, and (c) maximum expression value. These values were then merged across all normal tissue to generate comprehensive reference matrices: a junction-tissue matrix and an exon-tissue matrix. Cohorts with NAT samples (e.g., Cohort 1 and CPTAC) included NAT samples as part of the control tissue pool.

### Specific splicing junction detection module

Junctions were identified and quantified using the ASJA tool^[19]^, based on transcript assemblies. Junction expression was measured as coverage per ten million reads (CPT). When a junction was present in multiple transcripts, the transcript with the highest CPAT score and the highest transcript per million (TPM) value was selected as the representative isoform.

A splicing junction was classified as tumor-specific if it exhibited expression ≥5 CPT in tumor samples and met at least one of the following criteria relative to the junction-control tissue reference matrix:

1. Absolute novelty: completely absent from all control tissues;
2. Low background prevalence: detected in <1% of samples within any single control tissue and showing tumor expression at least 5-fold higher than the median value of any control tissue;
3. Robust tumor specificity: detected in <90% of samples from any control tissue, with tumor expression ≥5× the median and ≥10× the maximum value observed in any control tissue.

### Specific exon detection module

Exon identification and quantification were also performed using StringTie-assembled transcripts. Genomic coordinates and expression levels of individual exons were extracted from the assembled transcripts. Exon expression was normalized using the formula:e_ncov = (e_cov / unique mapped reads from STAR) × 10⁷, where e_cov refers to the exon coverage output from StringTie. If the number of uniquely mapped reads was unavailable, the sum of e_cov values for all known exons in the sample was used as an alternative denominator. For single-exon transcripts, genomic coordinates were adjusted based on their open reading frame (ORF) to enable appropriate comparison with controls. When an exon was present in multiple transcripts, the transcript with the highest TPM was selected as its representative.

Tumor-specific exons were identified using the same criteria applied to the exon-control tissue reference matrix, with an additional requirement that the exon expression exceeded 8 (e_ncov). To account for the limited resolution of untranslated region (UTR) annotations in RNA-seq data, we applied specific rules for comparing the first and last exons: For first exons, comparability was established only when the donor splice site was shared between tumor and control; For last exons, comparability required a shared acceptor splice site.

### TST-derived neoantigens prediction module

To predict TST-derived neoantigens, TSTs generated from tumor-specific splicing junctions and exons were first merged and filtered based on coding potential, as determined by both CPC2^[47]^ (v3.0.5) and CPAT^[48]^ (v0.1). Only those transcripts classified as coding by both tools and containing a complete open reading frame (ORF) were retained as coding TSTs (see *Transcripts coding potential prediction* section). To minimize false positives, only ORFs of coding TSTs directly resulting from tumor-specific transcriptional events (tumor-specific splicing junction and exon) were translated into protein sequences (in silico translation) by Biopython (v1.81). Potential novel protein sequence were identified by extracting 11-amino-acid windows upstream of each tumor-specific splicing junction or exon boundary, extended to encompass the full coding region. In cases where the tumor-specific transcriptional event occurred at or near the transcription start site (e.g., first exon or first junction), the entire translated protein sequence was subjected to peptide extraction to expand the immunogenic screening range. HLA class I genotyping was performed using arcasHLA^[49]^ (v3.9) for samples from Cohort 1, Cohort 2, and the CPTAC cohort. HLA typing results for TCGA-PAAD samples were obtained from the GDC (see *Prediction HLA Typing* section). Only HLA alleles supported by NetMHCpan were retained. In total, high-confidence HLA genotypes were determined for 401 samples.

NetMHCpan^[50]^ (v4.1) was used to predict 8-11 amino acid peptides binding to HLA, using the parameters ‘-f -inptype 0 -BA -xls -a’. Peptides with predicted binding affinity scores (IC50) <500 nM were classified as either strong binders (SB) or weak binders (WB) and considered candidate neoantigens. Final TST-derived neoantigens were defined as peptides absent in reference database.

### Comparative analysis of tumor-specific splicing junctions using SNAF

To evaluate the tumor specificity of splicing junctions identified by NeoAPP, we employed SNAF as a comparative tool. The analysis environment was configured using Docker (image: frankligy123/altanalyze:0.7.0.1) and applied to the breast and colorectal cancer sample datasets(GSE77661). In accordance with the SNAF official guidelines, in addition to the full set of GTEx samples, we incorporated the TCGA skin cancer cohort and GTEx skin tissues as additional controls. Tumor-specific splicing junctions were identified using the ‘snaf.initialize’ and ‘snaf.JunctionCountMatrixQuery’ functions under default parameters. Results were saved in the ‘NeoJunction_statistics_maxmin’ file. Junctions with the values “True” in the columns ‘cond’, ‘cond_add_tcga_control’, and ‘cond_add_gtex_skin’ were extracted for downstream analysis.

### Proteomic and Immunopeptidomes data analysis

Coding transcripts of each PDAC tumor sample were translated into protein sequences using Biopython (v1.81) to generate sample-specific protein databases. Protein sequences were then merged and deduplicated based on the ‘Metadata ID’ from the experimental design (downloaded from https://proteomic.datacommons.cancer.gov/pdc/study/PDC000270) to create custom protein and decoy databases with the philosopher database --custom command.

The spectrum files corresponding to each ‘Metadata ID’ were analyzed using matched custom protein and decoy databases. Raw mass spectrometry data were converted to mzML format using ThermoRawFileParser (v1.4.2). Searches were performed using MSFragger^[51]^ (v3.8) and the Philosopher^[52]^ (v5.0) workflow with the following parameters: precursor mass tolerance of ±10, fragment mass tolerance of ±20 ppm, and enzyme cleavage settings defined in the closed_fragger configuration file provided by MSFragger. Methionine oxidation (+15.994915) and serine TMT labeling (+229.162932) were specified as variable modifications, while cysteine carbamidomethylation (+57.021464) and lysine TMT labeling (+229.162932) were set as fixed modifications. Proteins were filtered with Philosopher using an FDR threshold of <0.01.

PDAC neoTSTs were combined, their protein sequences deduplicated, and then integrated with the human UniProt database to generate a custom reference for searching PDAC organoid mass spectrometry data (MSV000096853). Raw data were converted to mzML format and analyzed using Comet software within the MSFragger^[51]^ (v3.8) and Philosopher^[52]^ (v5.0). Precursor tolerance was set to ±10 ppm, and fragment ion m/z tolerance was set to ±10 ppm. Peptides ranging from 8 to 11 amino acids were searched using “unspecific” digestion parameters. Peptides and protein were filtered with Philosopher using an FDR threshold of <0.05.

### Cell Cultures

Mouse pancreatic tumor cells Panc02 (Cell Bank of Shanghai, Chinese Academy of Sciences, Strain No. SCSP-5468, RRID:CVCL_D627) were cultured in RPMI 1640 complete medium. Murine splenocytes were cultured in RPMI 1640 medium with GM-CSF (Peprotech, 315-03-20UG) added to 4 ng/ml. All complete medium were supplemented with 10% fetal bovine serum (BDBIO), penicillin (100 U/ml), and streptomycin (100 μg/ml). All of these cell lines were cultured at 37 ℃ with humidified 5% CO2.

### Representative neoTSTs Selection and Experimental Validation

A subset of representative F-neoTSTs and single-exon neoTSTs, was selected from PDAC samples for experimental validation. *In vivo* immunogenicity testing was performed in female B6-hHLA-A11:01/hB2M (Strain No. T064359) and B6-hHLA-A02:01/hB2M (Strain No. T064344) transgenic mice (8-10 weeks old, GemPharmatech, China). Coding sequences of selected neoTSTs containing predicted neoantigens were synthesized, cloned into a Takara mRNA expression vector, and polyadenylated to generate mRNA templates. These were used for in vitro transcription (IVT) with N¹-methylpseudouridine modification using the T7 High Yield RNA Synthesis Kit (Yeasen, 10633ES60). To formulate lipid nanoparticle (LNP)-encapsulated mRNA vaccines (mRNA-LNP), SM-102 ionizable lipid, cholesterol, DSPC, and DMG-PEG2000 were dissolved in ethanol and mixed with mRNA in 100 mM citrate buffer (pH 4.0) at a 3:1 volume ratio (ethanol:aqueous phases) using the INano™ L microfluidic mixer (Micro&Nano). After concentration and purification, the resulting mRNA-LNP formulations were obtained. Ten PDAC neoTSTs were randomly divided into two pools (5 µg mRNA each) and formulated into two mRNA-LNP vaccines. Humanized HLA-A02:01/HLA-A11:01 mice received intramuscular (IM) injections at two separate anatomical sites during the first and second weeks, respectively. In the third week, mice were euthanized, and splenocytes were isolated for immunological assessment by IFN-γ ELISpot and flow cytometry (see Supplementary Information).

To evaluate the anti-tumor efficacy of neoTSTs in the Panc02 syngeneic tumor model, representative neoTSTs and mutation-derived neoantigens were identified from Panc02 transcriptomic data (see Supplementary Information). neoTSTs and mutation neoantigens were formulated into separate mRNA-LNP vaccines using the method described above. For tumor induction, 3×10⁶ Panc02 cells were subcutaneously injected into the right flank of male C57BL/6N mice (n = 6 per group) on day 0. Mice received two IM doses of mRNA-LNP vaccine (5 µg mRNA per dose) on days 5 and 12. Tumor length (L) and width (W) were measured twice weekly, and tumor volume (V) was calculated as: V = L×W^2^ /2. On day 35, all mice were euthanized and splenocytes were collected for flow cytometry analysis.

### Additional information of computational and experimental methods

Detailed descriptions of the computational analyses (such as splicing junction annotation, scRNA-seq analysis, mutation-derived neoantigen prediction, representative neoTST selection, and other related methods), as well as experimental analysis for target validation (including cell culture, mRNA template synthesis, in vitro transcription, lipid nanoparticle formulation, flow cytometry, IFN-γ ELISpot assays, and related techniques), are provided in the Supplementary Information.

### Statistical analyses

Statistical analyses were performed using R (v4.0.2) and GraphPad Prism 8.0. Comparisons between two groups were assessed by the Wilcoxon rank-sum test. Survival analyses utilized Kaplan-Meier estimators with log-rank tests to determine statistical significance. One-way ANOVA was employed for group comparisons in tumor size and flow cytometry data; results are presented as mean±SEM. Significance levels were defined as follows: *P < 0.05, **P < 0.01, ***P < 0.001.

## Abbreviations

TSTs: Tumor-specific transcripts
neoTSTs: neoantigen-encoding TSTs
TSS: Transcription Start Site
EVs: Extracellular Vesicles
PDAC: Pancreatic ductal adenocarcinoma
TE: Transposable Element
ORF: Open Reading Frame

## Ethics approval and consent to participate

The clinical samples and information used in the study was approved by the Fudan University Shanghai Cancer Center (approval no.050432-4-1212B). The animal experiments were conducted in compliance with protocols approved by the Shanghai Medical Experimental Animal Care Commission.

## Consent for publication

Not applicable.

## Availability of data and materials

All data supporting this study are publicly accessible through the following repositories: RNA-seq BAM files and whole-exome sequencing (WES) VCF files for the CPTAC PDAC cohort were obtained from the GDC Data Portal (https://portal.gdc.cancer.gov/) using gdc-client; proteomic raw data (Accession: PDC000270) was retrieved from the Proteomic Data Commons (PDC); clinical annotations for the CPTAC cohort were sourced from Cao et al. (DOI: 10.1016/j.cell.2021.08.023); TCGA clinical metadata was acquired from Thorsson et al. (DOI: 10.1016/j.immuni.2018.03.023) with HLA typing data from the Pan-Cancer Atlas (https://gdc.cancer.gov/about-data/publications/panimmune); somatic mutation annotations (MAF files) were downloaded from the TCGA-PAAD repository (https://portal.gdc.cancer.gov/); somatic mutation-derived neoantigens (SNVs) for TCGA were obtained from TSNAdb (https://pgx.zju.edu.cn/tsnadb/download/); ChIP-seq BED files for transcription factors in pancreatic cell lines were acquired from ChIP-Atlas; PDAC single-cell RNA-seq data corresponds to GEO accession GSE212966; and RepeatMasker annotations (hg38) were sourced from the UCSC Genome Browser. Other data supporting the findings of this study are available from the corresponding author upon reasonable request.

## Acknowledgements

This work was supported by the National Key Research and Development Project of China (2021YFA1300500), and National Natural Science Foundation of China (82272625).

## Authors’ contributions

Conceptualization: S.H, Z.C; Methodology: S.H, J.Z; Visualization: J.Z,Y.Y, Q.L; Validation: Y.Y, Q.L; Software: P.L, Y.W; Funding acquisition: S.H; Project administration: S.H; Supervision:W.H, W.Y, Y.L, Z.H; Writing original draft: S.H, J.Z; Writing review & editing: S.H, J.Z,Q.L. All authors read and approved the final manuscript.

